# HetF protein is a new divisome component in a filamentous and developmental cyanobacterium

**DOI:** 10.1101/2020.09.09.290692

**Authors:** Wei-Yue Xing, Jing Liu, Zi-Qian Wang, Ju-Yuan Zhang, Xiaoli Zeng, Yiling Yang, Cheng-Cai Zhang

**Author notes:** Cheng-Cai Zhang,. Author Contributions J.-Y.Z., and C.-C.Z designed research; W.-Y.X., J.L, and Z.-Q.W performed research; X.Z., and C.-C.Z analyzed the data; W.-Y.X., Y.Y., and C.-C.Z. wrote the paper. **Statement of conflict of interests:** The authors declare no conflict of interests.

## Abstract

Bacterial cell division, with a few exceptions, is driven by FtsZ through a treadmilling mechanism to remodel and constrict the rigid peptidoglycan (PG) layer. Yet, different organisms may differ in the composition of the cell-division complex (divisome). In the filamentous cyanobacterium *Anabaena* PCC 7120, *hetF* is required for the initiation of the differentiation of heterocysts, cells specialized in N_2_-fixing cells under combined nitrogen deprivation. In this study, we demonstrate that *hetF* is expressed in vegetative cells and necessary for cell division in a conditional manner. Under non-permissive conditions, cells of a Δ*hetF* mutant stop dividing, consistent with increased level of HetF under similar conditions in the wild type. Furthermore, HetF is a membrane protein located at midcell and cell-cell junctions. In the absence of HetF, FtsZ rings are still present in the elongated cells; however, PG remodelling is abolished. This phenotype is similar to that observed with the inhibition of septal PG synthase FtsI. We further reveal that HetF is recruited to or stabilized at the divisome by interacting with FtsI and this interaction is necessary for HetF function in cell division. Our results indicate that HetF is a member of the divisome, and reveal distinct features of the cell-division machinery in cyanobacteria that are of high ecological and environmental importance.

**Significance Statement:** Cyanobacteria shaped the Earth’s evolutionary history, and are still playing important roles for elementary cycles in different environments. They are consisted of highly diverse species with different cell shape, size and morphology. Although these properties are strongly affected by the process of cytokinesis, the mechanism remains largely unexplored. Using different approaches, we demonstrate that HetF is a new component of the cell division machinery in the filamentous cyanobacterium *Anabaena* PCC 7120. The common and diverged characteristics of cell division in prokaryotes reflect the evolutionary history of different bacteria, as an adaptive measure to proliferate under certain environmental conditions. As a protein for cell differentiation, the recruitment of HetF to the septum illustrates such an adaptive mechanism for cyanobacteria.

## Introduction

Bacterial cell division is a highly complex and coordinated event that has been extensively studied in model bacteria (1, 2). In *Escherichia coli*, cell division starts with the polymerization of a tubulin-like protein FtsZ at midcell position, forming a membrane-associated structure called Z-ring that is stabilized and anchored to the inner cell membrane by proteins such as FtsA and ZipA (3, 4). This early Z-ring complex then serves as a scaffold for sequential recruitment of other cell division proteins, such as FtsK, FtsQ, FtsL, FtsB, FtsW and FtsN that are involved in chromosome segregation, structural integrity of the protein complex or transfer of precursors required for cell wall synthesis (5, 6). Then, more cell-division proteins related to PG synthesis (such as FtsI, PBP1b and LpoB) or hydrolysis (such as AmiA, AmiB and AmiC) are assembled into the structure, forming a large cell division complex, the divisome, that further executes the septum formation (6). When cell constriction starts, FtsZ guides PG remodeling at the outer surface of the plasma membrane through a treadmilling mechanism (7), which leads to the separation of the two daughter cells.

Although many components of the divisome are conserved in bacteria, some proteins of the machinery diverge significantly in subgroups of bacteria (8–10). In cyanobacteria, FtsZ is also conserved, so are many proteins involved in structural stability of the divisome and cell wall synthesis or chromosome segregation, such as FtsI (PBP3), PBP1b, AmiA, AmiB, AmiC, FtsK, FtsQ and FtsW. Other components, identified in cyanobacteria but not found in *E. coli*, are either shared by gram-positive bacteria such as *Bacillus subtilis*, or specific to cyanobacteria. For example, FtsA and ZipA are not conserved in cyanobacteria, but their functions are replaced by ZipN (Ftn2), ZipS (Ftn6) or SepF (Cdv2) (11–15). The composition of the cyanobacterial divisome has been investigated, and a network of interaction was revealed among many of these cell division related proteins (11–17). Despite all the effort, cell-division mechanism in cyanobacteria is still poorly understood in comparison to the model organisms such as *E. coli*.

As Gram-negative prokaryotes, cyanobacteria are ubiquitous and play essential roles in element cycles. Some cyanobacteria are able to differentiate into different cell types for division of labour. This is the case for *Anabaena* PCC 7120 (*Anabaena*) that is able to perform oxygen-labile N_2_ fixation using heterocysts, a specialized cell type regularly intercalated among vegetative cells which perform oxygen-evolving photosynthesis (18, 19). Heterocysts account for 5-10% of the total cells on the filaments and are induced upon deprivation of combined nitrogen in the growth medium. Heterocyst differentiation takes about 20-24 h, during which extensive morphological and physiological changes occur (18–20). Heterocyst development involves a complex regulatory network, among which NtcA and HetR are the central factors (21–24). Another regulator is HetF, which has been shown to be essential for heterocyst development in both *Nostoc punctiforme* ATCC 29133 and *Anabaena* (25, 26). HetF is a putative protease and mutation of the conserved active site led to a loss-of-function phenotype. Interestingly, beyond its function in cell differentiation, some reports show that *hetF* mutant displays morphological changes such as cell elongation and/or becoming larger in size (26), suggesting that *hetF* may also play a role in cell division. However, occurrence of such morphological changes was not consistently observed in the published data or even in the same study (25–28). Therefore, the function of *hetF* in cell division needs to be clarified.

Based on different approaches, we show here that HetF is involved in septal PG synthesis as a member of the divisome. These results advanced our understanding of the cell-division mechanism in cyanobacteria that are of ecological and environmental importance.

## Results

### *hetF* is expressed in vegetative cells and downregulated in mature heterocysts

*hetF* expression was described as constitutive but the experimental details were not provided (26). RNA-seq data indicates that the transcription start site (TSS) is located at -256 bp upstream of the *hetF* open-reading frame (29) (Fig. 1). *hetF* is preceded by an array of 12 small non-coding RNAs (*nsiR1*.*1 - nsiR1*.*12*) that are identical or highly similar to each other, all transcribed in the opposite direction of *hetF*. All the 12 copies of *nsiR1* have been shown to be activated early during heterocyst development following nitrogen step-down, in a NtcA and HetR-dependent manner (29–31). To investigate the cell-type specific expression of the *hetF* gene and the contribution of *nsiR1*, we made two *gfp* fusions with different promoters - a long version that covered the entire small RNA array (pP_*hetF*_*-gfp*) and a short version that included only the immediate surrounding region of the transcription start site (pP_*hetFa*_*-gfp*) (Fig. 1*A*). The replicative plasmid bearing each fusion was conjugated, respectively, into *Anabaena*. As shown in Fig. 1b, in a combined-nitrogen replete medium with nitrate (BG11), strong GFP fluorescence was detected in both strains without noticeable difference. Proheterocysts could be identified by alcian blue staining (32), we thus followed the changes of GFP fluorescence intensity during the process of heterocyst differentiation. At 15 hours post induction, the fluorescence level in the proheterocysts was either similar to or a little lower than that of vegetative cells. However, at 24-hour after induction, GFP fluorescence dropped to a low level in the mature heterocysts for both constructs. One copy of *nsiR1* (*nsiR1*.*1*) after the *hetF* TSS remained intact in pP_*hetFa*_*-gfp*; to inactivate this last copy, we either replaced the putative -10 box of *nsiR1*.*1* with a different sequence, or removed it from pP_*hetFa*_*-gfp*, which generated pP_*hetFb*_*-gfp* and pP_*hetFc*_*-gfp*, respectively. We found that these changes led to little variation of the GFP fluorescence under both combined-nitrogen replete or depleted conditions (Fig. 1). Our results indicate that *hetF* is highly transcribed in vegetative cells, but down-regulated in mature heterocysts. Such an expression pattern suggests a function of *hetF* in vegetative cells.

**Fig. 1.**
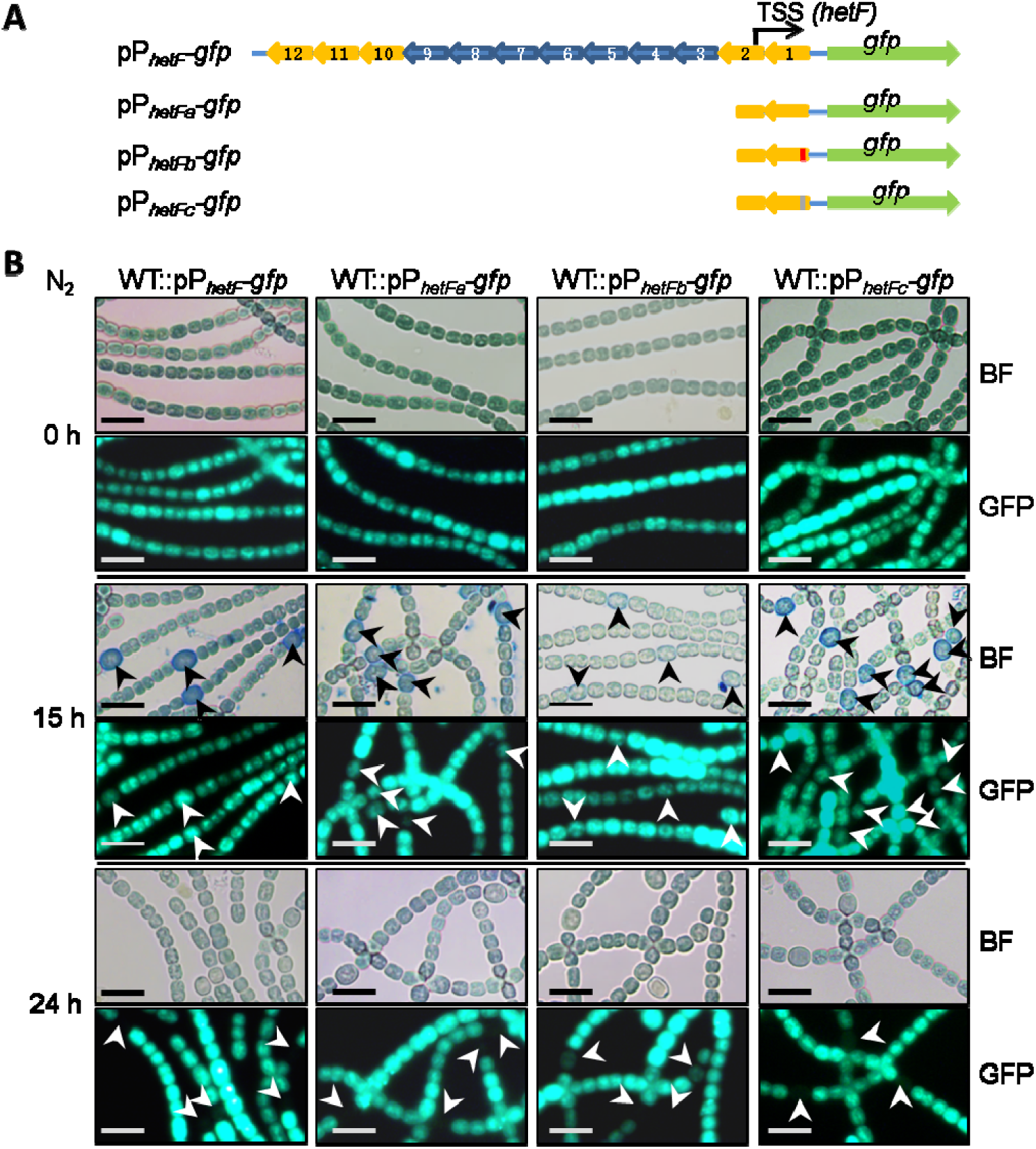
*hetF* is highly expressed in vegetative cells. (*A*) Sketch illustration of four transcription fusion constructs of *hetF*. The upstream *nsiR1* gene array is indicated by the blue and yellow arrows, genes with the same color are identical in sequence. The TSS (black arrows) of *hetF* locates in *nsiR1*.*2*. pP_*hetF*_-*gfp* includes the whole *nsiR1* array (*nsiR1*.*1 - nsiR1*.*12*). pP_*hetFa*_-*gfp* only includes *nsiR1*.*1*. pP_*hetFb*_-*gfp* and pP_*hetFc*_-*gfp* are generated from pP_*hetFa*_-*gfp* with the -10 box being replaced (black box) and deleted (gray box), respectively. (*B*) Expression pattern of *hetF* during heterocyst formation. The fluorescence pattern of WT cells harboring pP_*hetF*_-*gfp*, pP_*hetFa*_-*gfp*, pP_*hetFb*_-*gfp* or pP_*hetFc*_-*gfp* were captured after 0 h, 15 h and 24 h of nitrogen step-down (N_2_). BF: bright field. Cells were stained with alcian blue for better visualization of proheterocysts at 15 h. Arrows indicate proheterocysts or mature heterocysts. Scale bars, 10 μm.

### The cell-division defect of *hetF* mutant is dependent on light intensity and nitrogen regime

To investigate *hetF* functions, a markerless deletion mutant of *hetF* was created as described (33) (Fig. S1). As published (25, 26), we confirmed that the Δ*hetF* mutant could not form heterocysts, and consequently was unable to grow under diazotrophic conditions (Fig. S3). A dramatic cell-division defect, with cell elongation and slight enlargement, was observed under standard culture conditions for the mutant cells. However, this phenotype was not always reproducible, even for cells from the same culture. Thus, a continuous monitoring of the mutant phenotype was performed and the result showed that cell-division phenotype of Δ*hetF* changed throughout the growth course (Fig. 2*A*). With a fresh culture started at a low optical density (OD) around 0.08, cell filamentation appeared between day 2 and day 4, then the cell morphology returned to normal as the wild type (WT) with increasing OD values (Fig. 2*A*).

**Fig. 2.**
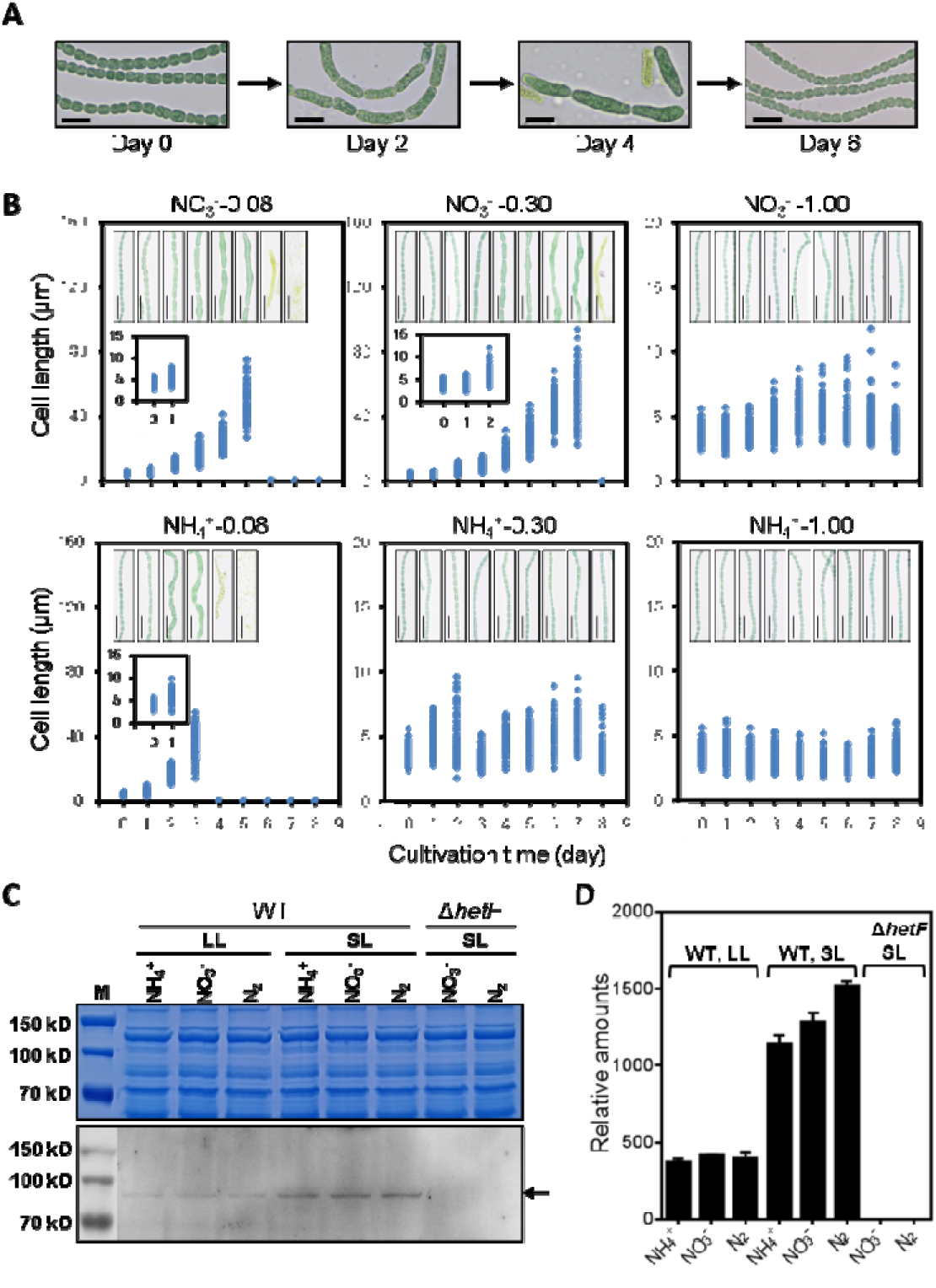
The cell filamentation phenotype of Δ*hetF* is dependent on light intensity and nitrogen regime. (*A*) Cell division defect of Δ*hetF* changes in the culture. Images of the Δ*hetF* cells taken at different day during cultivation in standard light (SL). Scale bars, 10 μm. (*B*) Effect of cell optical density (OD) and nitrogen sources on the mutant cell length. Continuous growth of the cell cultures at specified conditions (nitrogen source and the OD values were indicated on top of each panel) was achieved by daily dilution with respective fresh medium. 100 cells were measured for each sample and the representative microscopic images were presented (all cultures were grown in SL, please see more details in Methods). Zoomed-in plots were shown in some panels for better visualization of the data. Note that cell lysis occurred with time under conditions of low OD. Scale bars, 20 μm. (*C*) Western blot analysis of HetF expression level at day 2 of the different conditions corresponding to Fig. S2 using the multiclonal antibody against the CHAT domain (1-555aa) of HetF. (*D*) Quantification of HetF protein levels based on the western-blotting result from (*C*).

We sought to determine the factors accounting for the variation of cell-division phenotype of Δ*hetF*. Since cells at higher OD receive less light exposure due to the shading effect, we first investigated whether light intensity may be one of such factors. We thus followed the development of cell morphology with cultures maintained at low, medium and high OD respectively, with nitrate or ammonium as the nitrogen source, and with daily dilution of the cultures back to the initial OD value. As shown in Fig. 2*B*, in the culture with nitrate, cells at a low OD of 0.08 quickly developed cell-division defect as early as day 2, and cells continuously elongated, reaching 27 µm – 75 µm in length, while WT cells had a cell length ranging from 1.6 µm to 4.2 µm (Fig. 2*B* and Fig. S2*A*). From day 6, extensive cell lysis occurred and the culture collapsed, thus indicating that *hetF* was essential under such conditions. A similar process was found for cells maintained at a medium OD of 0.3, with slightly slower progression of the phenotype. However, cells grown and maintained at a high OD of 1.0 remained with a normal cell shape as that of WT (Fig.2*B*, upper panel); however, when exposed to a higher light intensity (HL), these cells could develop again the cell-division defect (Fig. S2*C*, left panel). These results indicated that increasing the illumination arrested cell division in the mutant even at high OD values. As a control, the WT cultured under similar light regimes showed little variation in cell length (Fig. S2 *A* and *B*).

In addition to light intensity, ammonium appeared to affect the phenotype as well, although less strongly. When ammonium was present as the nitrogen source, cells maintained at a low OD (0.08) showed a defect in cell division, but no significant cell filamentation was observed at medium OD (0.30) or high OD (1.00). Therefore, ammonium somehow suppressed the cell-division phenotype with increasing OD values (Fig.2*B*, lower panel). The effect of light and ammonium was unrelated to cell growth rate, because ammonium could support a faster cell growth than nitrate, but nevertheless led to a much weaker phenotype in cell division. Furthermore, we monitored the phenotype with cultures under low light (LL) or standard light (SL) with different nitrogen source. Our results show that most of the cells grown under LL with ammonium as the nitrogen source have a normal cell shape (Fig. S3), while conditions under SL with nitrate induced a cell-division-defect phenotype (Fig. S3).

Together, these results demonstrate that the phenotype of Δ*hetF* is dependent on light intensity and nitrogen regime, providing a rational for the inconsistency in the cell-division phenotype of the mutant in the published data (25-28). This study also allowed us to optimize the culture conditions, so that the function of *hetF* in cell division could be investigated consistently. For subsequent experiments, the *hetF* mutant was maintained under LL with ammonium where the cell could divide almost normally as the WT (thus defined as permissive conditions), and then shifted towards SL with nitrate at a low OD (non-permissive conditions) for phenotypic dissection.

Next, to better understand the conditional requirement of *hetF* in cell division, the level of HetF was examined by immunoblotting assay in the WT. The results showed that HetF level was about three-fold higher under SL than under LL condition regardless the nitrogen source used (Fig. 2 *C* and *D*). Therefore, the level of HetF increased in the cells under conditions where HetF is necessary for cell division.

### HetF localizes at midcell and cell-cell junctions

Many proteins involved in cell division display a midcell localization. We therefore examined the subcellular localization of HetF with the help of a GFP fusion. Initially, GFP sequence was fused to either the N- or C-terminus of HetF with a flexible peptide linker. However, neither of the fusions gave rise to detectable fluorescence. HetF is a multi-domain protein (Fig. 3*A*). We sought to minimize the interference of GFP folding in the fusion product. An analysis of the secondary structure using PSIPRED suggested the presence of two flexible regions in HetF, one from Q385 to C470 and another from C485 to W557. We therefore choose these two regions for making in-frame insertion of *gfp*, at 1275 bp and 1635 bp (corresponding to D425 and L545, respectively) within the coding region of *hetF*. When the fusions were expressed in *Anabaena*, only the one with GFP sequence inserted after D425 (HetF_D425_GFP, corresponding to the insertion of *gfp* at 1275 bp) showed fluorescence (see below). This fusion was functional because a construct expressing HetF_D425_GFP under the control of the native promoter P_*hetFa*_ complemented Δ*hetF* for both cell division (Fig. 3*B*) and heterocyst development (Fig. S4). This complemented strain allowed us to check the subcellular localization of HetF_D425_GFP by comparing with a WT filament pictured in the same frame under the miscroscope. HetF was mainly localized in the middle of cells just before physical constriction became visible (red arrows), or cell-cell junctions (yellow arrows). This localization pattern was similar to that observed after HADA labelling that marked PG synthesis at the septa (34).

**Fig. 3.**
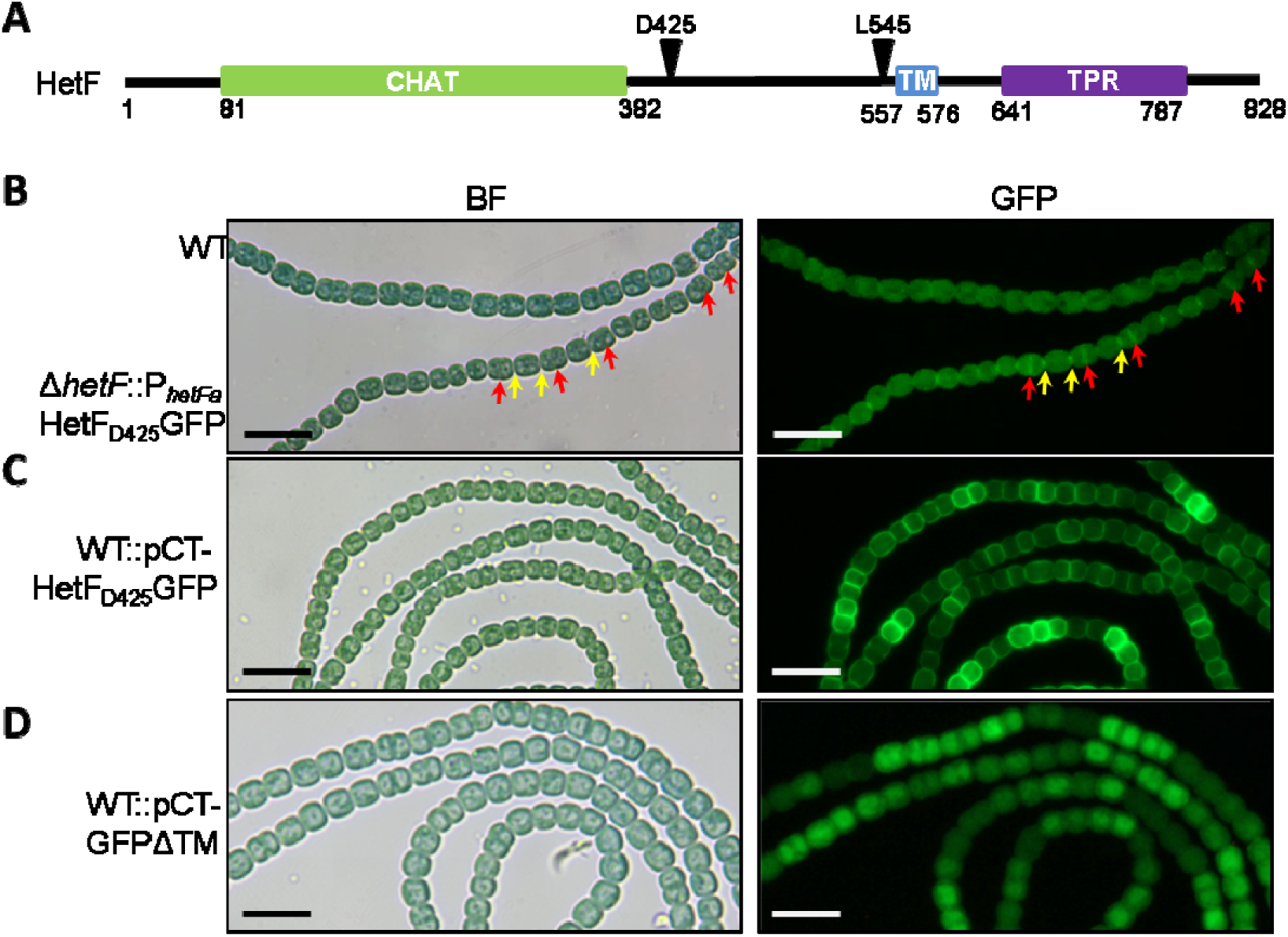
Subcellular localization of HetF in *Anabaena*. (*A*) Domain organization of HetF. GFP insertion sites are indicated by arrows. (*B*) subcellular localization of HetF_D425_GFP. Micrographs taken with a mixed culture of Δ*hetF*::pP_*hetFa*_-HetF_D425_GFP and WT strains grown under LL in BG11 (NO_3_^-^) medium. Red and yellow arrows indicate the localization of HetF_D425_GFP at midcell and cell-cell junctions, respectively. (*C*) and (*D*) HetF localizes on the cell membrane. In WT cells, HetF_D425_GFP expressed under CT promoter shows membrane localization (*C*), and removal of the transmembrane domain (GFPHetFΔTM) caused delocalization of the protein into cytoplasm (*D*). WT::pCT-HetF_D425_GFP and WT::pCT-GFPHetFΔTM were cultivated with inducers (4 mM theorphylline + 2 μM CuSO_4_) for 2 days under SL in BG11 (NO_3_^-^). BF: bright field. Scale bars in (*B*-*D*), 10 μm.

HetF is predicted to contain one trans-membrane (TM) segment (Fig. 3*A*). Consistent with this prediction, HetF_D425_GFP fluorescence was found mostly at the periphery of the cells when overexpressed under the control of a synthetic inducible promoter (CT) (33–35) (Fig. 3*C*). When the TM domain was deleted, the fluorescence was evenly distributed in the cytoplasm (Fig. 3*D*). We conclude therefore that HetF is a membrane protein localized at midcell and cell-cell junctions, consistent with the cell-division phentotype of Δ*hetF*.

### HetF acts at the step of septal PG synthesis

During cell division, septal PG synthesis is driven by FtsZ through a treadmilling mechanism (7). First, we examined whether the deletion of *hetF* affected the localization of FtsZ. To do so, the native form of *ftsZ* was first replaced by a *ftsZ-cfp* fusion on the chromosome (WT::*ftsZ-cfp*) and then *hetF* was deleted on the chromosome by homologous recombination (Δ*hetF*::*ftsZ-cfp*). When the Δ*hetF*::*ftsZ-cfp* strain was cultured under LL in the presence of ammonium, a few cells on the filaments were slightly elongated as compared to WT::*ftsZ-cfp*, but in most cells, FtsZ-CFP fluorescence was still found at midcell similar as in WT::*ftsZ-cfp* (Fig. 4*A*). When Δ*hetF*::*ftsZ-cfp* was cultured under SL with nitrate, extensive cell filamentation took place, a sign of cell-division arrest; under such non-permissive conditions, the Z-ring was still able to form, as one to three FtsZ-CFP ring-like structures were found in each elongated cell (Fig. 4*A*). Therefore, deletion of *hetF* led to cell filamentation but did not affect FtsZ localization.

**Fig. 4.**
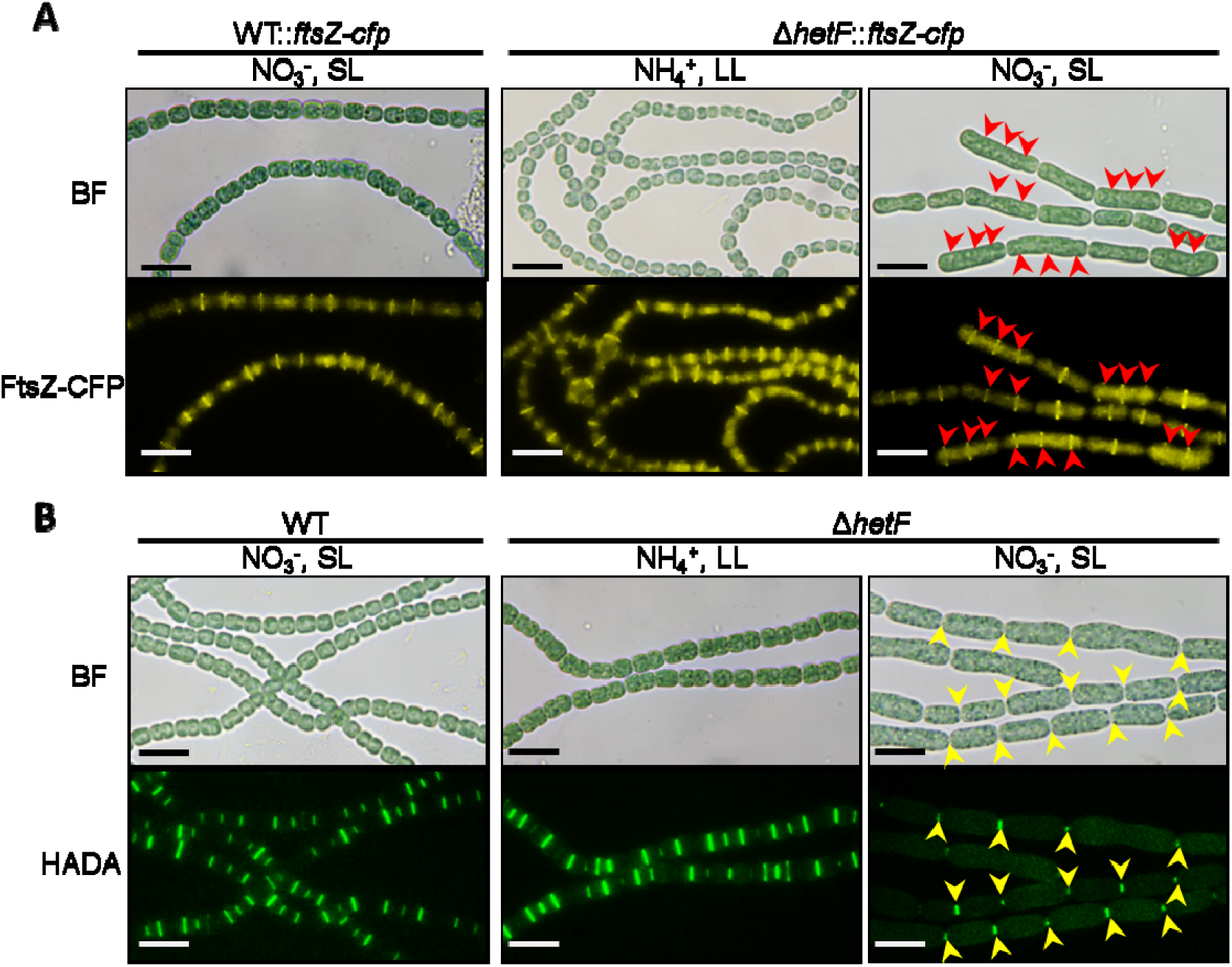
FtsZ-ring formation and HADA labelling of PG synthesis. (*A*) FtsZ ring formation in WT and the Δ*hetF* strain. The cell division defect of Δ*hetF*::*ftsZ-cfp* was induced by growing in BG11 (NO_3_^-^) under SL for 2 days. Red arrows indicate the FtsZ-CFP rings in the elongated cells. (*B*) PG synthesis indicated by HADA labelling. Cells from (*B*) were incubated with 200 µM of HADA for 1 day under SL or 2 days under LL, using nitrate or ammonium as nitrogen source as indicated. Yellow arrows indicate HADA fluorescence in the elongated cells. Scale bars, 10 μm. Pseudocolor for fluorescence was used and the original microscopic images were shown in Fig. S5.

Next, we checked the effect of *hetF* deletion on the later steps of cell division, namely septal PG remodelling. HADA is a fluorescence analogue of D-Ala that can be used to mark the site of septal PG synthesis in *Anabaena* filaments (34, 36). In WT cultured either under LL with ammonium (Fig. S6), or under SL with nitrate, cell division occurred normally and HADA fluorescence was observed at midcell or cell-cell junctions (Fig. 4*B*, left column). Similar HADA fluorescence pattern was found when Δ*hetF* was grown under LL with ammonium. However, when the mutant was transferred to non-permissive conditions, no HADA labelling could be found along the elongated cells, except those at the cell-cell junctions that represented the cell poles from the precedent cell-division cycle (Fig. 4*B*, middle and right column). This observation suggested that although FtsZ rings were still formed in the elongated cells, HADA incorporation, and hence PG remodelling, did not take place in the absence of HetF. Taken together, our results show that HetF is required for cell division at the step of septal PG synthesis.

### HetF interacts with FtsI and this interaction is required for cell division

Localization of HetF at midcell position and its requirement for septal PG synthesis suggest that HetF could be directly involved in cell division as a component of the divisome. As a major enzyme for PG synthesis in the divisome, FtsI/PBP3 is known to be a target of the antibiotic aztreonam, which inhibits septal cell-wall synthesis (37). Previously, we found that impairment of FtsI by aztreonam led to cell filamentation and abolished PG synthesis at septal sites, but did not affect FtsZ-ring formation (34). This observed effect of aztreonam was therefore similar to that of *hetF* deletion, and was further confirmed in this study (Fig. 4 and Fig. S7). We thus tested whether HetF could interact with FtsI. Using a bacterial two-hybrid system (BACTH), we showed that HetF indeed interacted with FtsI when either of the partners was expressed on the two different vectors of BACTH (Fig. 5*A*).

**Fig. 5.**
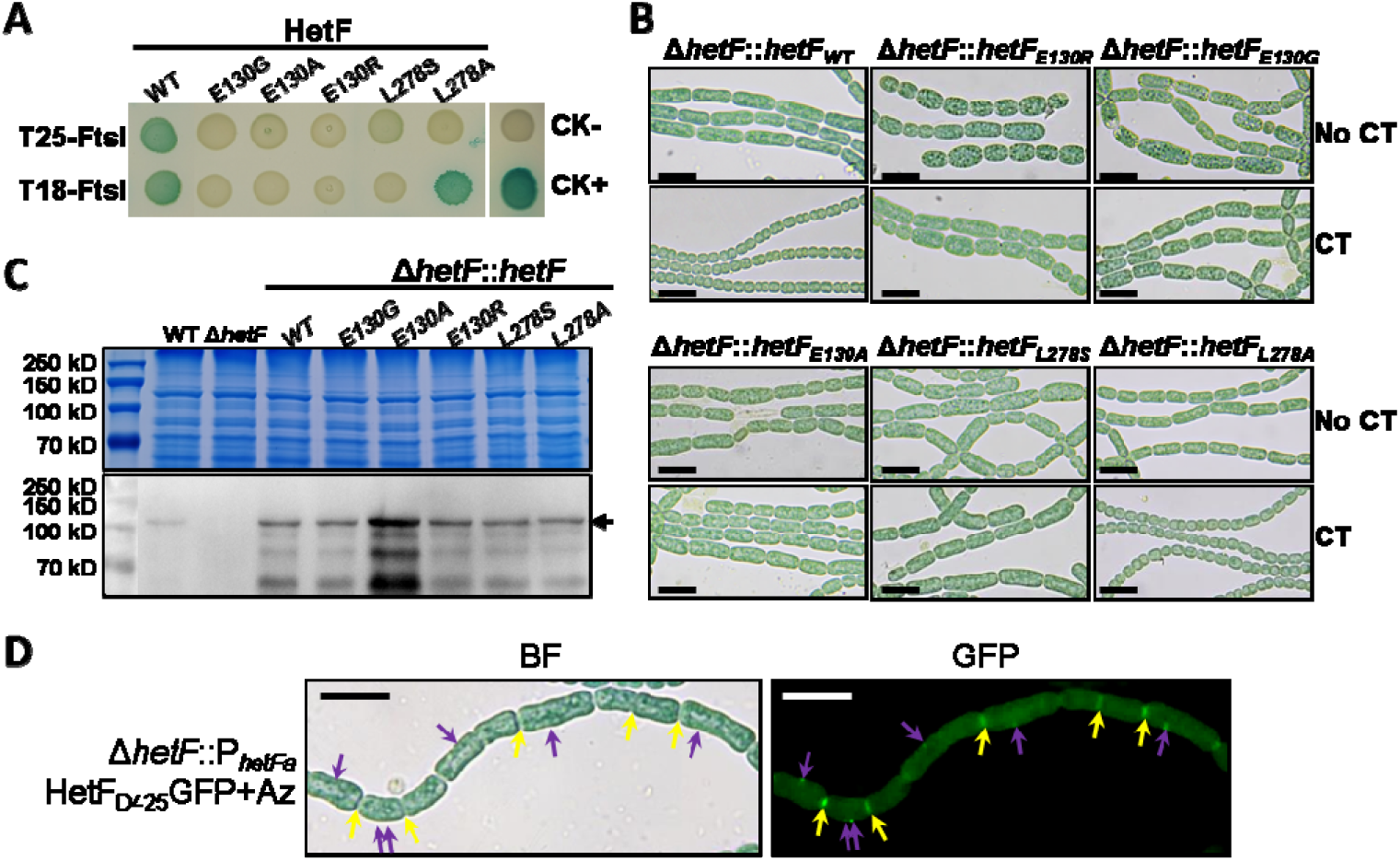
Interaction between HetF and FtsI is required for cell division. (*A*) Binary protein-protein interaction detected by the BACTH assays for HetF (WT) or its variants with point mutation expressed using either T25 or T18 vector. Positive interaction between two targeted proteins is indicated by the blue colour of the colony, and negative interaction by white ones (see details in Methods section). CK- and CK+ indicate negative and positive controls of BACTH assays, respectively. (*B*) Mutant HetF proteins with point mutations unable to interact with FtsI failed to complement the cell division defect of Δ*hetF* (please see details in Methods section), in contrast to the WT form of HetF. CT, copper and theophylin for induction of *hetF* expression. (*C*) Western-blot analysis of HetF levels in the strains from (*B*) using HetF antibody. Total protein extracts (upper panel) using for western blotting (lower panel) were used as controls for protein amounts. (*D*) Micrographs of Δ*hetF*::pP_*hetF*_-HetF_D425_GFP after 72-h cultivation in BG11 treated with 100 μM aztreonam under LL. Yellow arrows indicate the HetF_D425_GFP subcellular localization at cell-cell junctions. Purple arrows indicate the HetF_D425_GFP foci at the side wall of cells. BF: bright field. Scale bars in *B* and *D*, 10 μm.

To further test the specificity and function of FtsI-HetF interaction, we used BACTH to screen mutations that rendered HetF unable to interact with FtsI. After screening from a *hetF* mutant library generated by error-prone PCR, two point mutations of HetF, E130G and L278S, were identified, and HetF with either of the two mutations lost the ability to interact with FtsI (Fig. 5*A*). Further substitution of E130 with Ala or Arg (E130A and E130R) gave the same results, demonstrating that the E130 residue is crucial for the interaction of HetF with FtsI. On the other hand, when L278 of HetF was replaced by Ala residue, this mutant form of HetF retained the ability to interact with FtsI, but only when HetF_L278A_ was expressed from the T18 vector. To test the effect of such mutations *in vivo*, WT *hetF* or its different mutant versions was placed behind the inducible CT promoter and used in complementation assays of Δ*hetF* in *Anabaena*. As a control with WT *hetF*, without induction by copper and theophylline (TP), the Δ*hetF* still showed defect in cell division, but this defect was restored after addition of inducers (Fig. 5*B*). In contrast, *hetF*_*E130R*_, *hetF*_*E130G*_, *hetF*_*E130A*_ or *hetF*_*L278S*_ failed to complement the cell-division defect of Δ*hetF* even after induction, in accordance with the inability of their corresponding proteins to interact with FtsI. Interestingly, *hetF*_*L278A*_ which could still encode a protein able to interact with FtsI, retained the ability to complement Δ*hetF* (Fig. 5*B*). To rule out that the mutations making *hetF* non-functional is caused by protein stability, we checked the protein levels using polyclonal antibodies raised against HetF with Western blotting (Fig. 5*C*). Similar amounts of total proteins were loaded on the gel for different strains (upper panel). HetF, with an apparent molecular weight of 91.8 kD, was detected in WT but not in Δ*hetF* (lower panel). In all strains, HetF mutant forms was detected at levels even higher than in the WT, which is expected since these proteins were expressed from a replicative plasmid pCT (33, 35). Therefore, the point mutations did not affect HetF stability. Together, these results demonstrate that the interaction of HetF with FtsI is essential for the function of HetF in cell division.

### Subcellular localization of HetF requires a functional FtsI

Since HetF is part of the divisome through interaction with FtsI, we examined whether a functional FtsI was required for the proper subcellular localization of HetF. For this purpose, the florescence of HetF_D425_GFP was examined in cells following the inhibition of FtsI by adding aztreonam (Fig. 5*D*). While the fluorescence of HetF_D425_GFP was still observed at cell-cell junctions, formed after the preceded cell cycle, in the elongated cells caused by inhibition with aztreonam, only small patches or foci of fluorescence could be observed. These results indicated that once the function of FtsI was inhibited, HetF_D425_GFP could no longer be localized or stablized at septa. Together with the interaction between HetF and FtsI, these results indicate that the recruitment of HetF to, or its stabilization at, the divisome is dependent on FtsI.

## Discussion

In this study, we provide evidence that HetF participates directly in cell division in *Anabaena* as a member of the divisome. This conclusion is supported by several lines of evidence. First, *hetF* is strongly expressed in vegetative cells under nitrogen-replete conditions (Fig. 1*B*). This expression pattern is different from those of genes critical for heterocyst development, such as *hetR, ntcA* or *patS*, whose expression is low in vegetative cells but increased in developing cells and mature heterocysts (21–24, 38). Instead, *hetF* expression profile is similar to those reported for genes involved in cell division, such as *ftsZ*, with a downregulation in mature hetercysts (37, 39), consistent with the terminal nature of this cell type. Secondly, Δ*hetF* displays a dramatic cell filamentation phenotype, a clear sign of cell-division defect. This phenotype is the strongest when the mutant was cultured under conditions of increasing illumination with nitrate as a nitrogen source, and cells maintained under such conditions lysed following cell elongation, indicating that *hetF* was essential under such conditions. Low light intensity, and to a lesser extent ammonium, could suppress this phenotype. Thirdly, HetF displays a midcell localization in cells ready to divide, and this localization persists till the end of the cell division at the cell-cell junctions as some other cell-division proteins and HADA labeling patterns in *Anabaena* (16, 34). In agreement with the subcellular localization, HetF interacts with FtsI as demonstrated by both BACTH and genetic evidence based on point mutations of HetF that disrupts FtsI-HetF interaction. Furthermore, the *hetF* deletion mutant and a WT strain treated with aztreonam to inhibit FtsI give comparable results for FtsZ localization and HADA labeling. Finally, the proper subcellular localization of HetF requires a functional FtsI. Thus, HetF is recruited to or stabilized at the septum through its interaction with FtsI.

Our data also clarified the inconsistency of cell division phentotypes of *hetF* mutant in previous work (25–28). Although it is still unknown how light and nitrogen source affect the function of HetF, we observed that under conditions where HetF is required for cell division, the protein levels of HetF increase (Fig. 2 *C* and *D*). On the other hand, the *hetF* mutant never differentiates heterocysts, whether the cell division defect appeared or not (Fig. S3). Thus, the conditional requirement of *hetF* is specific to its function in cell division, whereas its role in heterocyst development is essential.

Although major components involved in cell division are highly conserved in different bacteria, our study indicates that cyanobacteria have distinct features for their cell division machinery. The role of HetF in cell division may correspond to a mechanism for these organisms to adapt to the changing environment, such as the light intensity, which is not only the energy source but also a shaping force for the whole physiology of photosynthetic organisms. Beyond the common features in bacterial cell division, the evolutionary history of particular bacteria leads to divergence and variation in cell division for better environmental adaptation.

## Materials and Methods

The procedures for the construction of *Anabaena* mutant strains are described in detail in *SI*

### Strains and Growth conditions

All strains used in this study were listed in Supplementary Table S1. *Anabaena* strains were cultivated in BG11 (40) or BG11_0_ medium (BG11 without nitrate), in a shaker (30°C, 180 rpm) with standard light illumination of 30 μmol photons m^-2^s^-1^ (SL). When necessary, cultures were incubated at low light illumination of 7 μmol photons m^-2^s^-1^ (LL) or high light illumination of 70 μmol photons m^-2^s^-1^ (HL). In order to maintain a stable cell division arrest phenotype, the Δ*hetF* culture was cultivated and stored in BG11_0_ with 2.5 mM NH_4_Cl under LL. Cell-division-arrest phenotype could be induced by cultivating in BG11 under SL with an initial OD_750_ of 0.08 – 0.3 for 2-4 days. 100 μg mL^−1^ neomycin or 5 μg mL^−1^ spectinomycin and 2.5 μg mL^−1^ streptomycin were added to the cultures as needed. All strains are listed in Table S1.

### Assay for testing the effect of cell density, light intensity and nitrogen sources on cell division arrest

Δ*hetF* and WT were washed with BG11_0_ for three times to remove NH_4_Cl and resuspended in BG11_0_ with 2.5 mM NH_4_Cl or BG11 to the OD_750_ = 0.08, 0.3, and 1, respectively. The cultures were then incubated under SL or HL. The value of OD was measured every day and the culture was then washed and diluted to the respective initial OD value again using corresponding fresh medium. Meanwhile, samples were collected every day for recording the phenotypes. The cell length was averaged from 100 cells.

### Testing the cell division arrest phenotype in Δ*hetF* background

Δ*hetF* background strains were washed with BG11 for three times to remove NH_4_Cl before resuspended in BG11 to OD of 0.08 - 0.30. The cultures were incubated under SL for 2 days, then samples were prepared for observing the cell division arrest phenotype. For complementation experiment, the cell-division-arrest phenotype of Δ*hetF*::pCT-HetF was induced by cultivating in BG11 under SL for 2 days, then the cultures were diluted to OD of 0.08 - 0.30 in BG11 supplemented with 2 μmol CuSO_4_ and 4 mM theophylline, and continuously incubated under SL for 1-2 days before recording the cell phenotypes. The same method was used for the strains expressing *hetF* point mutations, such as Δ*hetF*::pCT-HetFE130R.

### Multiclonal antibody preparation

The plasmid pHTAlr3546CHATStrep was constructed by inserting a partial *hetF* fragment (1-1665 bp, amplified with primers Palr3546F1d / Palr3546R1665f) and a synthesized TwinStreptag fragment into pET28a. The expressed fusion protein contains the CHAT domain (1-555 aa) of HetF, a 6His-tag at its N-terminal and a Twin-strep tag at its C-terminal. To induce CHAT domain protein expression, *E. coli* BL21 (DE3) cells containing pHTAlr3546CHATStrep were grown in LB medium with 0.5 µM IPTG at 37°C. After 4 hours’ induction, cells were collected and resuspended in lysis buffer (pH 7.4, 137 mM NaCl, 2.7 mM KCl, 8 mM Na_2_HPO_4_, 14.6 mM KH_2_PO_4_) and were broken by JN mini-low temperature and ultrahigh pressure cell breaker (JNBIO Co. Ltd.). The lysate was centrifuged at 10000 rpm for 40 min at 4°C. The precipitate (Inclusion body) was washed twice with 2 M urea, and then sent to Frabio Co. Ltd. for multiclonal antibody production.

### Preparation of *Anabaena* total protein and Western blot assay

Cells were collected by filtration and then broken by a sample preparation system (FastPrep-24, 6.0 m/s, QuickPrep, 60 s) in LDS-sample loading buffer (70 mM Lithium dodecyl sulfate, 100 mM Tris-HCl (PH 8.5), 10 % Glycerol, 4 mM EDTA, 0.025 % Coomassie brilliant blue G250). Cell extracts were boiled at 95°C for 10 min followed by centrifugation at 135,000 rpm for 10 min. The supernatant was collected and loaded to 10% SDS-PAGE and subsequently subjected to western blot assay with a multiclonal antibody against the CHAT domain in HetF.

### BACTH assay

The Bacterial Adenylate Cyclase Two-Hybrid System Kit (BATCH) based on the reconstitution of adenylate cyclase was used for testing protein-protein interaction (41). *hetF* and *ftsI* were amplified with primers listed in Supplementary Table S2 and individual PCR product were assembled into linearized pUT18C and pKT25 vectors, respectively. All the resulting plasmids were verified by PCR and Sanger sequencing. The plasmids were co-transformed into strain BTH101, and the transformants were plated on solid LB medium containing 50 μg L^-1^ Apicillin, 25 μg L^-1^ Kanamycin, 0.5 mM L^-1^IPTG and 40 μg L^-1^ X-gal.

For Screening of HetF point mutations that lost interaction with FtsI, *hetF* ORF region was amplified using Green Taq Mix (Vazyme Biotech Co., Ltd, P131-AA) and primers (Palr3546F1 / Palr3546R2502) for 15 cycles. The PCR product was assembled into pKT25 vector using One Step Cloning Kit (Vazyme Biotech Co., Ltd, ClonExpress II). The ligation product was transformed into BTH101 containing T18-FtsI plasmid. White colonies appeared on the plates were selected for colony PCR and check the presence of single missense mutations of HetF via Sanger sequencing.

### Microscopy

The SDPTOP EX30 microscope was used to take brightfield images and the SDPTOP EX40 for fluorescent images. HADA fluorescence were excited with 405 nm and captured at 460 nm, similar to the blue fluorescent protein (36). The filter (EX379-401, DM420LP, EM435-485) was used to image HADA fluorescence (Exposure time: 200 ms). The filter (EX426-446, DM455LP, EM460-500) was used to image CFP fluorescence (Exposure time: 1 s). The filter (EX470-490, DM495LP, EM500-520) was used to image GFP fluorescence (Exposure time: 1 s). All images were processed using ImageJ.

## Supporting information

Supplemental Materials with additional methods, figures and tables.

## Acknowledgments

This work was supported by the Key Research Program of Frontier Sciences of the Chinese Academy of Sciences (Grant No. QYZDJ-SSW-SMC016).

