## Supplemental Materials with additional methods, figures and tables. for "HetF protein is a new divisome component in a filamentous and developmental cyanobacterium"

Cheng-Cai Zhang

.

**This PDF file includes:**

Supplementary text

Figures S1 to S7

Tables S1 to S3

SI References

Supplementary Information Text

**SI materials and methods**

**Construction of *Anabaena* mutant strains***. hetF* was constructed in a similar way as previously described using pCfp1 plasmid (1). The repair template was generated by fusing the upstream and downstream region of the target sequence using primers listed in Table S2. The spacer sequence was designed according to the described rules (1) and prepared by annealing two complementary primers. To generate the plasmid that knocks out a specific gene, the respective repair template and the spacer sequence were sequentially cloned into pCpf1 at the sites of *Bgl*II-*BamH*I and *Aar*I−*Aar*I.

To construct *gfp* transcriptional reporter strains for *hetF*, the promoter regions (−2276 to 30 for P***_hetF_***-*gfp* and −300 to 30 for P***_hetFa_***-gfp) of *hetF* were amplified with specific primers and subsequently cloned into *BamH*I-*Xho*I digested plasmid pRL25N-L*gfp* (2), resulting in plasmids pP***_hetF_***-*gfp* and pP***_hetFa_***-*gfp*. The plasmid pP***_hetFb_***-*gfp* is a derivative of pP***_hetFa_***-*gfp* bearing a mutation in the -10 box of *nsiR1*. This mutation is created via site-directed mutagenesis on pP***_hetFa_***-*gfp* using a pair of primers (Palr3546F166m and Palr3546R173m) that have the desired mutation in the overlapping sequence. pP***_hetFc_***-gfp was also derived from pP***_hetFa_***-*gfp,* through deletion of the -10 box of *nsiR1* using the same strategy.

To overexpress HetF using a CT promoter, the ORF region of *hetF* (1 to 2484 with respect to the start codon) amplified using the primers Palr3546F1f and Palr3546R2484e, was cloned into pCT at the sites of *Xho*I-*Sma*I (1, 3), resulting in the overexpression plasmid pCT-HetF. Plasmids bearing HetF variants with point mutations, pCT-HetFE130R, pCT-HetFE130G, pCT-HetFE130A, pCT-HetFL278S and pCT-HetFL278A, were constructed by site-directed mutagenesis using pCT-HetF as the template and respective primer pairs containing desired mutation (Palr3546E130R-seqF / Palr3546E130R-seqR, Palr3546E130G-seqF / Palr3546E130G-seqR, Palr3546E130A-seqF / Palr3546E130A-seqR, Palr3546L278S-seqF / Palr3546L278S-seqR, and Palr3546L278A-seqF / Palr3546L278A-seqR, respectively).

To check HetF localization in *Anabaena*, two translational fusions under different promoters were constructed using a similar strategy as described above. For the plasmid with native promoter, the repair template was generated by fusing 3 fragments (−300 to 1275 bp relative to *hetF* coding region, *gfp*, and 1276 to 2484 bp of *hetF*) by overlapping PCR. The repair template was sequentially cloned into pCT at the sites of *BamH*I-*Sma*I (1, 3), resulting in the translational fusion plasmid pP*_hetF_*-HetF_D425_GFP. The plasmid with CT promoter (for over-expression) was constructed in the same way, but using a repair template without the native promoter sequence (1 to 1275 bp of *hetF*, *gfp*, and 1276 to 2484 bp of *hetF*). The repair template was cloned into pCT at the sites of *Xho*I-*Sma*I (1, 3), resulting in the translational fusion plasmid pCT- HetF_D425_GFP.

Plasmid pCT-GFPHetFΔTM was used for checking the localization of a HetF variant without the TM domain*.* The repair template was generated by fusing 3 fragments (*gfp*, 1 to 1665 bp of *hetF*, 1732 to 2484 bp of *hetF*) using overlapping PCR. The repair template was cloned into pCT at the site of *Xho*I-*Sma*I (1, 3), resulting in the translational fusion plasmid pCT-GFPHetFΔTM.

Plasmid pRLAlr3858-CFP for checking the FtsZ localization in *Anabaena* was constructed with the suicide vector pRL277 (2). The repair template was generated by fusing 4 fragments (161 to 1284 bp of *ftsZ*, *cfp*, Km-resistant cassette, 1353 to 2375 bp of *ftsZ* coding region) using overlapping PCR. The repair template was then cloned into *Bgl*II-*Xho*I cutted pRL277 (2), resulting in the translational fusion plasmid pRLAlr3858-CFP.

To make the *Anabaena* mutant strains, the constructed plasmid (see Table S3) was transferred into *Anabaena* by conjugation (4, 5). The exconjugants were selected on BG11 plates containing appropriate antibiotics and subsequently verified by PCR and Sanger sequencing.

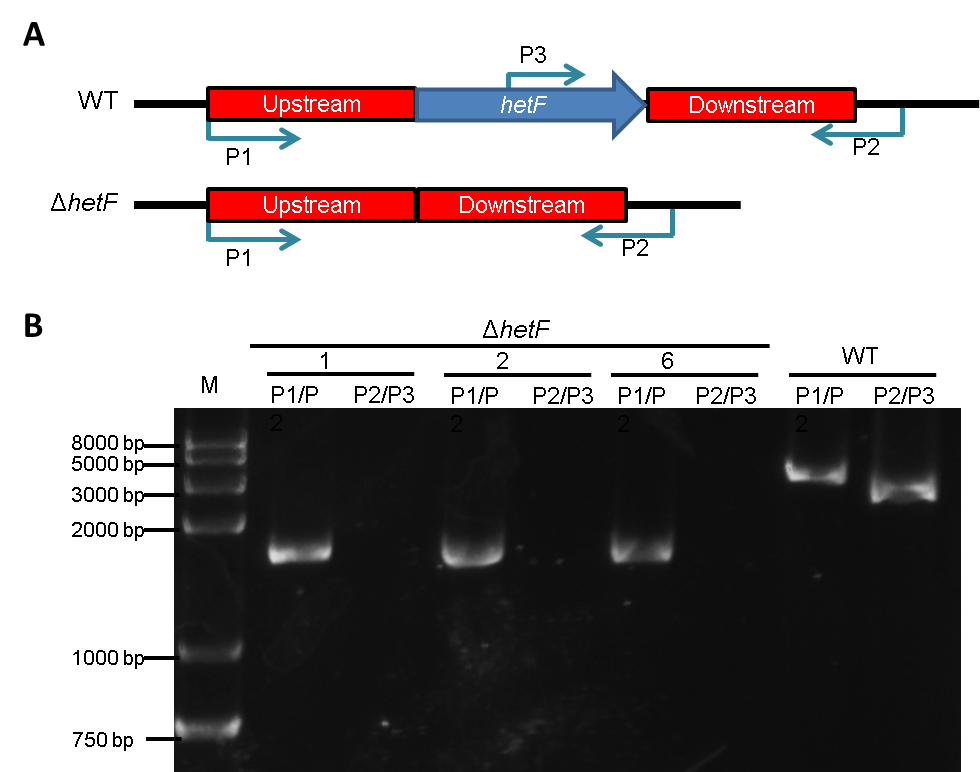

Fig. S1. Deletion of *hetF* and verification of Δ*hetF* genotype. (*A*) Schematic representation of the genotypes of WT and Δ*hetF*. Upstream: upstream homologous arm, Downstream: downstream homologous arm. The mutant was constructed by homologous recombination using the Cpf1 gene editing technique (see Materials and methods). (*B*) PCR Verification of the mutant genotype using the primers shown in panel A (blue arrows). P1, P2 and P3 were short names for the oligonucleotides Palr3546F300m, Palr3546R3435 and Palr3546E130A-seqF, respectively.

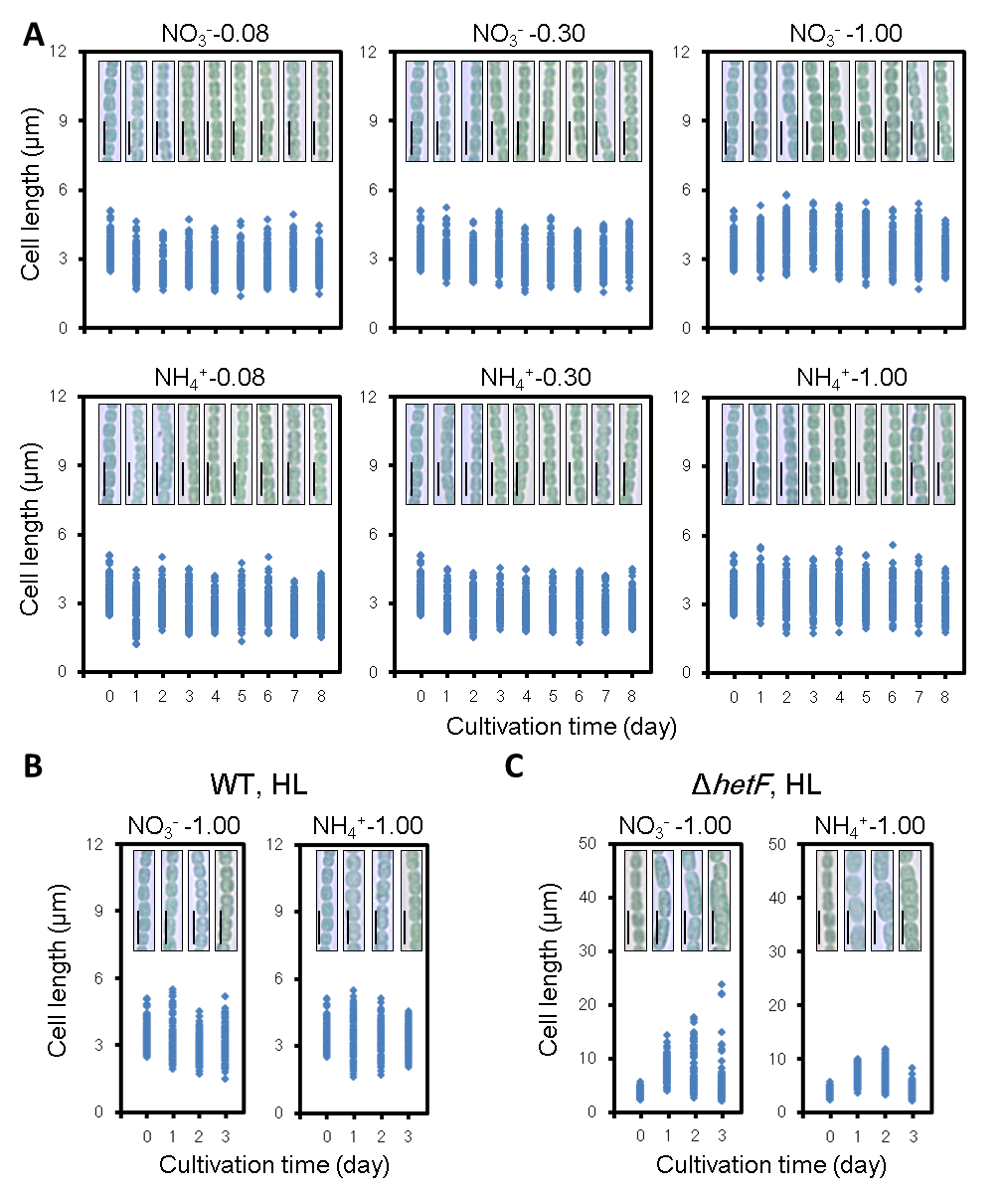

Fig. S2. Effect of the culture OD, light intensity and nitrogen sources on WT and Δ*hetF* cell length. (*A*) Cell length changes of WT strain grown under indicated conditions. These experiments were performed and the data were quantified as that described in Fig. 2. *B* and *C*, Effect of high light (HL) and nitrogen sources on the cell length of WT (*B*) and Δ*hetF* stain (*C*). Scale bars, 5 μm. The nitrogen source and the OD value of respective culture were indicated on the top of each panel..

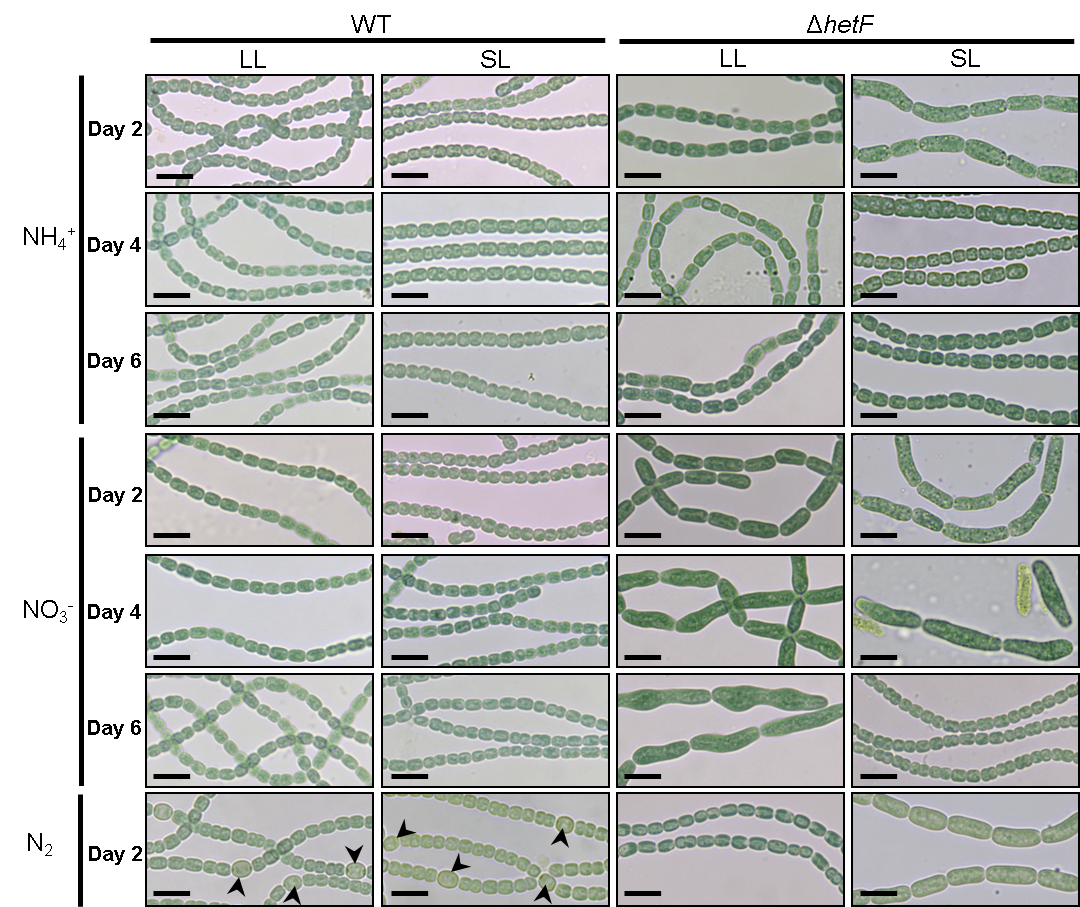

Fig. S3. The phenotype of cell division of *hetF* mutant is dependent on the culture conditions. With a starting OD_750_ of 0.08, images of WT and Δ*hetF* cells were taken at day 2, day 4 and day 6, respectively, during cultivation under different light intensity (LL or SL) and nitrogen sources (NH_4_^+^ or NO_3_^-^ ). To test the differentiation of heterocysts, samples of day 2 using NH4^+^ as nitrogen source were taken, and transferred into combined-nitrogen-free medium (N_2_) and incubated further for 2 days, followed by observation under a microscope. Note that although cells in the culture from LL with ammonium have little phenotypes in cell division, no heterocysts were induced. Arrows indicate heterocysts. Scale bars, 10 μm.

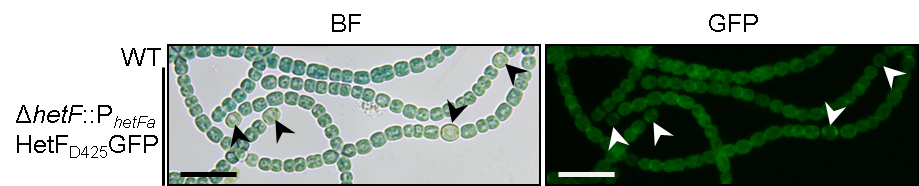

**Fig. S4.** Heterocyst differentiation of Δ*hetF*::pP*_hetFa_*-HetF_D425_GFP after 2 days cultivation under LL in BG11_0_ (N_2_). Arrows indicated heterocysts. BF: bright field. Scale bars, 10 μm.

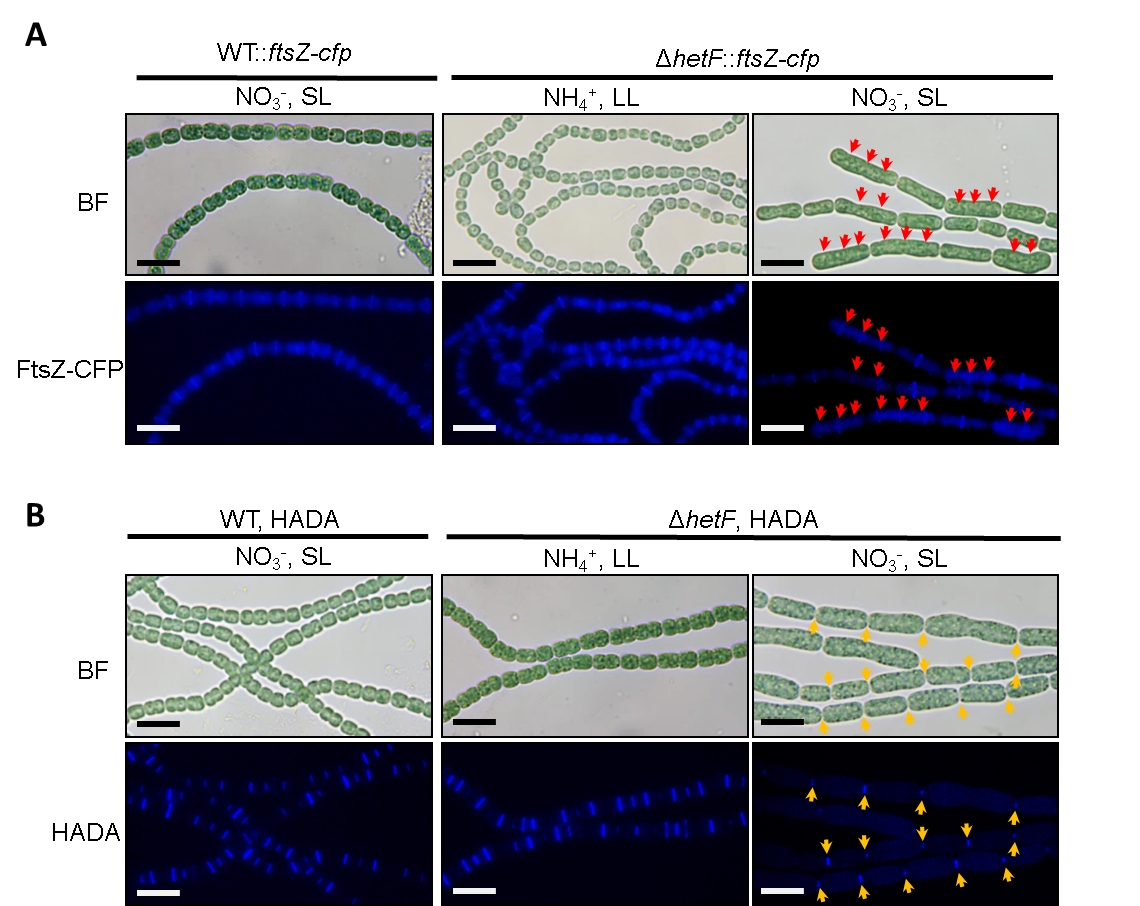

Fig. S5. Original images of Fig. 4. Please see the figure in main text for the legend.

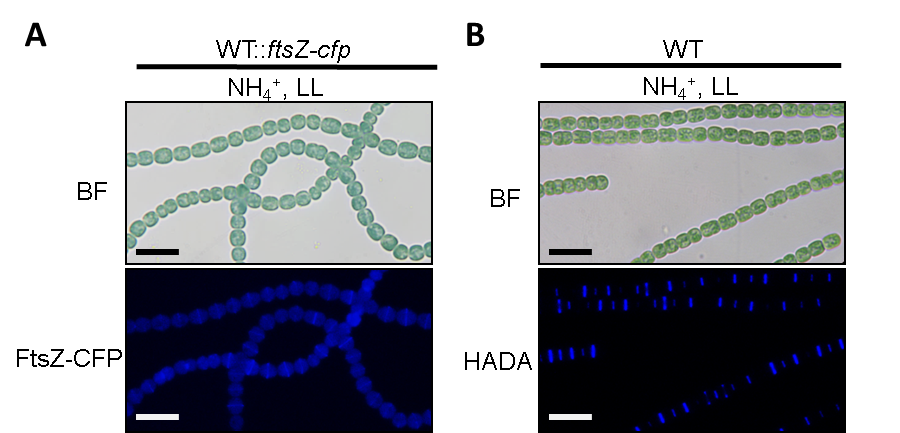

Fig. S6. FtsZ-ring formation (*A*) and HADA labelling of PG synthesis (*B*) in WT under LL using ammonium as the nitrogen source. For HADA labeling, cells were incubated with 200 µM of HADA for 2 days before imaging. Scale bars, 10 μm.

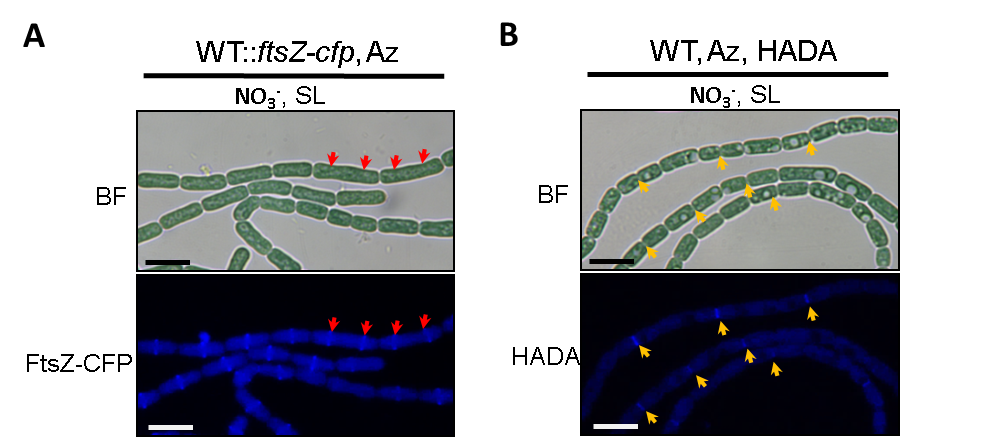

Fig. S7. FtsZ-CFP subcellular localization and HADA labelling after addition of the antibiotic aztreonam targeted to FtsI. (*A*) Micrographs of WT::*ftsZ-cfp* with aztreonam treatment. The strain WT::*ftsZ-cfp* was cultivated in BG11 (NO_3_^-^) with 100 μM aztreonam for 2 days under SL. Red arrows indicate the FtsZ-CFP rings in two representative elongated cells. (*B*) Micrographs of WT labelled with HADA after aztreonam treatment. WT was cultivated in BG11 with 100 μM aztreonam for 1 day under SL, then 200 µM of HADA was added and continuously cultivated for 1 day. Red arrows indicate HADA fluorescence at cell-cell junctions. BF: bright field. Scale bars, 10 μm.

Table S1. Strains used in this study.

| Strains | Description | Source |
| --- | --- | --- |
| *Anabaena* PCC 7120 | Wild type | Pasteur Culture Collection |
| WT::pP*_hetF_*-gfp | Wild type bearing pP*_hetF_*-gfp; Nm^r^ | This study |
| WT::pP*_hetFa_*-gfp | Wild type bearing pP*_hetFa_*-gfp; Nm^r^ | This study |
| WT::pP*_hetFb_*-gfp | Wild type bearing pP*_hetFb_*-gfp; Nm^r^ | This study |
| WT::pP*_hetFc_*-gfp | Wild type bearing pP*_hetFc_*-gfp; Nm^r^ | This study |
| Δ*hetF* | *hetF* markless delition mutant | This study |
| Δ*hetF*::pCT-HetF | Δ*hetF* bearing pCT-HetF; Nm^r^ | This study |
| Δ*hetF*::pCT-HetFE130R | Δ*hetF* bearing pCT-HetFE130R; Nm^r^ | This study |
| Δ*hetF*::pCT-HetFE130G | Δ*hetF* bearing pCT-HetFE130G; Nm^r^ | This study |
| Δ*hetF*::pCT-HetFE130A | Δ*hetF* bearing pCT-HetFE130A; Nm^r^ | This study |
| Δ*hetF*::pCT-HetFL278S | Δ*hetF* bearing pCT-HetFL278S; Nm^r^ | This study |
| Δ*hetF*::pCT-HetFL278A | Δ*hetF* bearing pCT-HetFL278A; Nm^r^ | This study |
| WT::*ftsZ-cfp* | *ftsZ* translational fusion with *cfp* on chromosome; Nm^r^ | This study |
| Δ*hetF*::f*tsZ-cfp* | *hetF* markless delition mutant and *ftsZ* translational fusion with *cfp* on chromosome; Nm^r^ | This study |
| Δ*hetF*::pP*_hetF_*-HetF_D425_GFP | Δ*hetF* bearing pP*_hetF_*-HetF_D425_GFP; Nm^r^ | This study |
| Δ*hetF*::pCT- HetF_D425_GFP | Δ*hetF* bearing pCT-HetF_D425_GFP; Nm^r^ | This study |
| Δ*hetF*::pCT-GFPalr3546ΔTM | Δ*hetF* bearing pCT-GFPalr3546ΔTM; Nm^r^ | This study |

Table S2. Primers used in this study.

| Primer | Sequence (5’-3’) |
| --- | --- |
| Palr3546F300m | GCAGAAATTCGATATCTAGATCGGGAATTGGTAAATTTTCCCTG |
| Palr3546R194 | AGCGCTACCGACGCTAGTGCT GAAAGAAACGCATCTCCCTGT |
| Palr3546F2501d | AGCACTAGCGTCGGTAGCGCT GAGAGATTGGGGAGTGGAGA |
| Palr3546R3382 | CGCAACGTTGTTGCCATTGCTGAGTAATTGCTGAGTGTAGTCA |
| cr_alr3546R848F | AGATACCAACAAACCCGCTAAATCGT |
| cr_alr3546R848R | AGACACGATTTAGCGGGTTTGTTGGT |
| Palr3546F2276m | ACATGGATCCTGTAGCTCTGTGTCTCTTG |
| Palr3546R30 | CACATCTCGAGGGTTACAGAAATGTGAAATTCC |
| Palr3546F300ma | ACATGGATCCGGGAATTGGTAAATTTTCCCTG |
| Palr3546F166m | CGTTAAACACAAGAGGGCAGATGCTAACCAATC |
| Palr3546R173m | CTGCCCTCTTGTGTTTAACGGTAGATGCACCTTGA |
| Palr3546F163m | AAACCCCCAGTGGCTCAGATGCTAACCAATCCGGATAA |
| Palr3546R221m | GCATCTGAGCCACTGGGGGTTTTTTCTATAACTAGCA |
| Palr3546F300mc | CGGCGGGGTTTTTTTTTGGA GGGAATTGGTAAATTTTCCCTG |
| Palr3546R2484e | ACCACCAGAACCCCC CTTGGGGCTTTTTTGTTGCAGA |
| Palr3546F1276c | GTGGTAGCACTAGCGTCGGT GATGGGGAAATGTCTTTACCGAT |
| Palr3546R1275c | GAGGCCTTGGATCCAGTCATATCCAGCACACCAGAGTAAGTAG |
| PYFP2-seF | ATGACTGGATCCAAGGCCTCT |
| PYFP2-seR | ACCGACGCTAGTGCTACC |
| Palr3546F1f | GAGGTAACAACAAGATGGTGTCCCAGGAATTTCACATT |
| Palr3546F1p | GTGGTAGCACTAGCGTCGGT GTGTCCCAGGAATTTCACATTTC |
| Palr3546R1665c | TGTCTGGCGACTTTGCTTGCGACGGACACGATT |
| Palr3546F1732b | CGTGTCCGTCGCAAGCAAAGTCGCCAGACAACAGTAG |
| Palr3546E130R-seqF | AGCACGTCTGCCGTGGCGAGTGATGCACGCAGGCGAT |
| Palr3546E130R-seqR | CGCCTGCGTGCATCACTCGCCACGGCAGACGTGCTAAC |
| Palr3546E130G-seqF | AGCACGTCTGCCGTGGGG AGTGATGCACGCAGGCGAT |
| Palr3546E130G-seqR | CGCCTGCGTGCATCACTC CCCACGGCAGACGTGCTAAC |
| Palr3546E130A-seqF | AGCACGTCTGCCGTGGGCAGTGATGCACGCAGGCGAT |
| Palr3546E130A-seqR | CGCCTGCGTGCATCACTGCCCACGGCAGACGTGCTAAC |
| Palr3546L278S-seqF | CCTCTGTGGCGACGATTCAGCGGGTTTGTTGGTTAAC |
| Palr3546L278S-seqR | TAACCAACAAACCCGCTGAATCGTCGCCACAGAGGGTT |
| Palr3546L278A-seqF | CCTCTGTGGCGACGATGCAGCGGGTTTGTTGGTTAAC |
| Palr3546L278A-seqR | TAACCAACAAACCCGCTGCATCGTCGCCACAGAGGGTT |
| Palr3546R1665f | GCTACCACCACCAGAACC CTTGCGACGGACACGATT |
| Palr3546F1d | TGCCGCGCGGCAGCCAT GTGTCCCAGGAATTTCACATT |
| Palr3858F161 | GCAGAAATTCGATATCTAGATCGTATTGGCGAGATTGTTCCTGG |
| Palr3858R1284 | TCCACCAGAGGCCTTGGATCCATTTTTGGGTGGTCGCCGTC |
| Palr3858F1353 | ACCGGATCATCAGTACTCCCTGCTAATTTTCAAGTTCAGAGGT |
| Palr3858R2375 | CGCAACGTTGTTGCCATTGCAGGTAGAACTTGTACCAGTGCA |
| PgfpspF | GGATCCAAGGCCTCTGGTGGATCTG |
| PgfpspRa | GGGAGTACTGATGATCCGGTGATT |
| Palr0718F1c | ATCACCTCTAGTGGTGAAATGCAAAAGTCACCAAGTAGAT |
| Palr0718R1830 | GATGTCGATCTAGATCTCTTAAGGTTTTCCTTCAATCGGCT |
| Palr3546F1 | ATCACCTCTAGTGGTGAAGTGTCCCAGGAATTTCACATT |
| Palr3546R2502 | GATGTCGATCTAGATCTCTCATCTTCCCGTACTCTACTT |

Table S3. Plasmids used in this study.

| Plasmid | Description | Source |
| --- | --- | --- |
| pCT | Km^r^Nm^r^; | 1, 3 |
| pCpf1-sp | Sm^r^ Sp^r^; CRISPR-Cpf1-Based Genome Editing vector | 1 |
| pCpf1 | Km^r^ Nm^r^; CRISPR-Cpf1-Based Genome Editing vector | 1 |
| pSfgfp-Sp | Sm^r^ Sp^r^; carrying supper foldding *gfp* coding sequence | 2 |
| pRL277 | Km^r^Nm^r^; | 2 |
| pBAD-mTurquoise2 | Km^r^Nm^r^; carrying cfp coding sequence | Addgene number:http://www.  addgene.org/54844/ |
| pRL25N-Lgfp | Km^r^ Nm^r^; carrying *gfp* coding sequence | 2 |
| pCpf1-alr3546R848-sp | Sm^r^ Sp^r^; CRISPR-Cpf1 editing plasmid for *hetF* markless delition | This study |
| pP*_hetF_*-gfp | Km^r^ Nm^r^; pRL25N-Lgfp carrying *hetF* promoter ( -2276 to 30 with respect to the start codon of hetF) and *gfp* coding gene | This study |
| pP*_hetFa_*-gfp | Km^r^ Nm^r^; pRL25N-Lgfp carrying *hetF* promoter ( -300 to 30 with respect to the start codon of hetF) and *gfp* coding gene | This study |
| pP*_hetFb_*-gfp | Km^r^ Nm^r^; the - 10 box of *nsiR1* that existed in *hetF* promoter based on pP*_hetFa_*-gfp had been mutant | This study |
| pP*_hetFc_*-gfp | Km^r^ Nm^r^; the - 10 box of *nsiR1* that existed in *hetF* promoter based on pP*_hetFa_*-gfp had been deleted | This study |
| pCT-HetF | Km^r^ Nm^r^; pCT carrying *hetF* ORF and CT promoter, used for over-expression | This study |
| pCT-HetFE130R | Km^r^ Nm^r^; pCT carrying *hetF* ORF with E130R point mutation and CT promoter, used for over-expression | This study |
| pCT-HetFE130G | Km^r^ Nm^r^; pCT carrying *hetF* ORF with E130G point mutation and CT promoter, used for over-expression | This study |
| pCT-HetFE130A | Km^r^ Nm^r^; pCT carrying *hetF* ORF with E130A point mutation and CT promoter, used for over-expression | This study |
| pCT-HetFL278S | Km^r^ Nm^r^; pCT carrying *hetF* ORF with L278S point mutation and CT promoter, used for over-expression | This study |
| pCT-HetFL278A | Km^r^ Nm^r^; pCT carrying *hetF* ORF with L278A point mutation and CT promoter, used for over-expression | This study |
| pP*_hetF_*-HetF_D425_GFP | Km^r^ Nm^r^; pCT carrying *hetF* ORF with gfp fusion and native promoter, used for hetF localization; gfp was inserted after 1245 with respect to the start codon of *hetF* | This study |
| pCT- HetF_D425_GFP | Km^r^ Nm^r^; pCT carrying *hetF* ORF with gfp fusion and CT promoter, used for *hetF* localization and over-expression; gfp was inserted after 1245 with respect to the start codon of *hetF* | This study |
| pCT-GFPHetFΔTM | Km^r^ Nm^r^; pCT carrying *hetF* ORF without TM domain (1666-1731) with gfp fusion and CT promoter, used for ΔTM localization and over-expression; gfp was inserted at the N-terminal of ΔTM | This study |
| pHTHetFCHATStrep | Km^r^ Nm^r^; pHTwinStrep carrying *hetF* ORF (1-1665) , used for protein expression and purification | This study |
| pRLAlr3858-CFP | Km^r^ Nm^r^; pRL277 carrying 161 to 1284 of *ftsZ* *+ cfp + kanamycin resistance cassette +* 1353 to 2375 of *ftsZ*, used for FtsZ localization | This study |
